# Differential metabolic activity and discovery of therapeutic targets using summarized metabolic pathway models

**DOI:** 10.1101/367334

**Authors:** Cankut Çubuk, Marta R. Hidalgo, Alicia Amadoz, Kinza Rian, Francisco Salavert, Miguel A. Pujana, Francesca Mateo, Carmen Herranz, Jose Carbonell-Caballero, Joaquín Dopazo

## Abstract

**Background:** in spite of the increasing availability of genomic and transcriptomic data, there is still a gap between the detection of perturbations in gene expression and the understanding of their contribution to the molecular mechanisms that ultimately account for the phenotype studied. Alterations in the metabolism are behind the initiation and progression of many diseases, including cancer. The wealth of available knowledge on metabolic processes can therefore be used to derive mechanistic models that link gene expression perturbations to changes in metabolic activity that provide relevant clues on molecular mechanisms of disease and drug modes of action (MoA). In particular, pathway modules, which recapitulate the main aspects of metabolism, are especially suitable for this type of modeling.

**Results:** we present Metabolizer, a web-based application that offers an intuitive, easy-to-use interactive interface to analyze differences in pathway module metabolic activities that can also be used for class prediction and in silico prediction of Knock-Out (KO) effects. Moreover, Metabolizer can automatically predict the optimal KO intervention for restoring a diseased phenotype. We provide different types of validations of some of the predictions made by Metabolizer.

**Conclusions:** Metabolizer is a web tool that allows understanding molecular mechanisms of disease or the MoA of drugs within the context of the metabolism by using gene expression measurements. In addition, this tool automatically suggests potential therapeutic targets for individualized therapeutic interventions.

Metabolizer can be found at: http://metabolizer.babelomics.org.

## Background

Because of their multigenic nature, cancer and other complex diseases are often better understood as failures of functional modules caused by different combinations of perturbed gene activities rather than by the failure of a unique gene [1]. In fact, an increasing corpus of recent evidences suggest that the activity of well-defined functional modules, like pathways, provide better prediction of complex phenotypes, such as patient survival [2,3], drug effect [4], etc., than the activity of their constituent genes. In particular, the importance of metabolism in cancer [5] and other diseases [6] makes of metabolic pathways an essential asset to understand disease mechanisms and drug MoA and search for new therapeutic targets.

Gene expression changes have been used to understand pathway activity in different manners. Initially, conventional gene enrichment [7] and gene set enrichment analysis (GSEA) [8] were used to detect pathway activity from changes in gene expression profiles [9]. However, these methods provided an excessively simplistic view on the activity of complex functional modules that ignored the intricate network of relationships among their components. Some modifications of enrichment methods, specifically designed for signaling pathways, took into account the connections between genes [10]. Nevertheless, such approaches still produced a unique value for pathways that are multifunctional entities and did not take into account important aspects such as the integrity of the chain of events that triggers the cell functions. More recently, mechanistic models focuses into the elementary components of the pathways associated to functional responses of the cell [3,11], providing in this way a more accurate picture of the cell activity [12]. Specifically, in the context of metabolic pathways, Constraint Based Models (CBM) have been applied to find relationship between different aspects of the metabolism and the phenotype [13]. CBM using transcriptomic gene expression data allowed the analysis of human metabolism in different scenarios at an unprecedented level of complexity [14,15]. However, as many mathematical models, CBM present some problems, such as their dependence on initial conditions or the arbitrariness of some assumptions, along with difficulties of convergence to unique solutions [13,16]. Moreover, with limited exceptions [17], most of the software that implement CBM models only run in commercial platforms, such as MatLab and working with them require of skills beyond the experience of experimental researchers. In spite of the complexity of metabolism, metabolic modules have been defined to provide a comprehensive curated summary of the main aspects of metabolic activity and account for the production of the main classes of metabolites (nucleotides, carbohydrates, lipids and amino acids) [18]. Here we present a simple model that accounts for the activity of metabolic modules [18] taking into account the complex relationships among their components and the integrity of the chain of biochemical reactions that must occur to guarantee the transformation of simple to complex metabolites. The likelihood of such reactions to occur is inferred from gene expression values within the context of metabolic modules. In order to make these models accessible and easily usable to the biomedical community, we have developed Metabolizer, an interactive and intuitive web tool for the interpretation of the consequences that changes in gene expression levels within metabolic modules can have over cell metabolite production.

## Implementation

### Inferring the metabolic activity of a KEGG module

Pathway Modules [18] are used to depict the complex interactions among proteins carrying out the reactions that account for the main metabolic transformations in the cell. Here, a total of 95 modules were used, that comprise a total of 446 reactions and 553 genes (Additional Table 1). The pathway modules were downloaded through REST-style KEGG API from the KEGG MODULE (http://www.genome.jp/kegg/module.html) database in plain text format files that include information of the metabolites, genes and reactions. Metabolic pathways were downloaded from KEGG PATHWAY database in KGML format files. Then, each KEGG module is made up of reaction nodes (composed by one or several isoenzymes or enzymatic complexes [19]), which are connected by edges in a graph that describes the sequence of reactions that transforms simple metabolites into complex metabolites, or vice-versa. The potential catalytic activity level of a KEGG module can be derived from the potential catalytic activities of all the reaction nodes, assuming all the intermediate metabolites are present and available. Under this modeling framework, the potential for catalytic activity of a reaction node is inferred from the presence of the constituent proteins. However, given the difficulty of obtaining direct measurements of protein levels, an extensively used proxy for protein presence is the observation of the corresponding mRNA within the context of the module [3,11,20–24]. Then, the contribution of potential catalytic activities of all individual nodes to the whole module metabolic activity can be derived by using a recursive method based on the Dijkstra algorithm [25]. Assuming a value of 1 for the initial node of the module, the potential catalytic activity of the subsequent nodes is calculated by the formula:

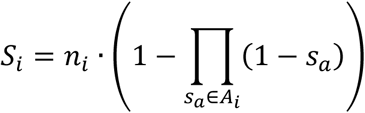

**Table 1.** TCGA samples used in this study

| Cancer Type | Abbreviation | Tumor samples | Normal Samples | Patients alive | Deceased patients |
| --- | --- | --- | --- | --- | --- |
| Breast Invasive Carcinoma | BRCA | 1057 | 113 | 900 | 146 |
| Kidney Renal Clear Cell Carcinoma | KIRC | 526 | 72 | 345 | 173 |
| Liver Hepatocellular Carcinoma | LIHC | 294 | 48 | 184 | 112 |
| Prostate Adenocarcinoma | PRAD | 379 | 52 | 367 | 7 |
| Total |  | 2256 | 285 | 1796<sup>1</sup> | 438<sup>1</sup> |

<sup>1</sup> The sum of these columns does not equal the total number of tumor samples plus normal samples because survival information was missing for some patients

In the formula, *S_i_* is the catalytic activity of the current node *i, n_i_* is the catalytic node activity value inferred from normalized gene expression values of the current node *i*, *A_i_* is the set of edges arriving to the node *i* that, within this modeling framework, accounts for the flux of metabolites produced by the corresponding reactions in other nodes with activity values *s_a_*.

The resulting integrity value of the whole sequence of reactions represented in the module is summarized by the value of catalytic activity propagated until the last node, which carries out the last the transformations of the chain of reactions that ultimately produces the final metabolite. This method is an adapted version of the propagation algorithm on graphs successfully used to estimate cell signaling activities in cancer [3]. Additional Figure 1 outlines the procedure.

**Figure 1.**
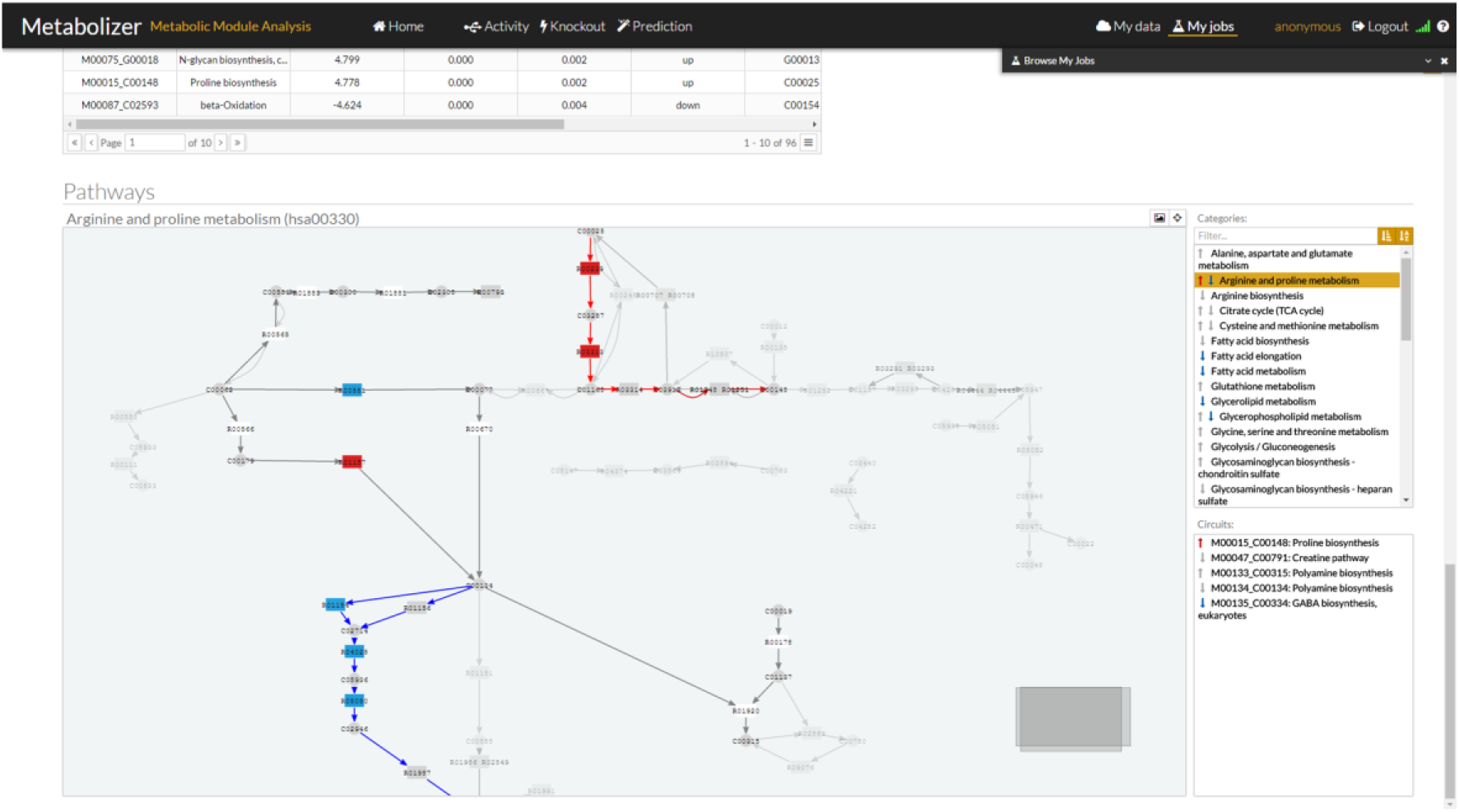
Metabolizer graphic interface with a representation of the modules. On the right side there is a list of KEGG pathways with arrows up or down in case they contain modules with up or down activations, respectively. When the arrow is gray, the change in activity is not significant. Red up arrows indicates a significant increase in activity and blue down arrow a significant decrease of activity in the module. Below the pathway list, there is another list with the modules within the pathway with the same code for arrows.

### Differential metabolic module activity estimation

Similarly to normalized gene expression values, module metabolic activity values calculated in this way make sense only within a comparison context, which allows deciding whether the estimated metabolic activity of a given module has changed significantly across the compared conditions or not. Here, Wilcoxon test is used to assess the significance of the observed changes of module metabolic activity when samples of two conditions are compared. Since many modules are simultaneously tested, multiple testing effects need to be corrected. FDR method [26] is used for this purpose.

### Class prediction

The class prediction functionality includes two sub-functionalities: training process, where the predictor is built using a training set, and classification process, where the predictor can be used for class prediction purposes.

In order to build a predictor a training set composed by samples belonging to two or more classes is required. The selection of samples that properly represent the variability of the classes is critical for the generalizability of the predictor. Two powerful prediction algorithms, Random Forests (RF) [27], as implemented in the R *randomForest* package (https://cran.r-project.org/web/packages/randomForest/), and Support Vector Machines (SVM) [28], as implemented in the R *e1071* package (https://cran.r-project.org/web/packages/e1071/), can be chosen to train the predictor. The algorithm uses the profiles of metabolic module activities of the two or more groups of samples compared. The accuracy obtained by the predictor is assessed by k-fold cross-validation and the area under the Receiver Operating Characteristic (ROC) curve.

Once a model has been trained, the predictor can be saved and can be used in a second phase to classify unknown samples. Thus, using the option “Test existing model” in the Metabolizer web interface, a list of samples can be uploaded and the proper predictor can be selected from the list of saved predictors. The predictor chosen will return a table with the probabilities of belonging to any of the classes for each sample.

### Prediction of the impact of KOs in metabolism

The model proposed can be used not only to derive metabolic module activity profiles in real conditions but also in simulated conditions. Therefore, KOs or over-expressions, alone or in combinations, can easily be simulated by changing the values of the targeted genes to 0 or 1 (or to any other low or high value between 0 and 1), respectively. Then, the simulated condition is compared to the original condition and a fold change threshold of 2 (that can be modified by the user) can be used to detect the most relevant changes in module metabolic activity. Since only two individual conditions (before and after KO) are compared a conventional test cannot be applied here.

In addition to individual gene interventions, the effect of drugs with known targets (as described in DrugBank [29]) over the different metabolic modules can be studied. It is possible to simulate the effect of drugs alone, in combinations, or combined with gene KOs or over-expressions. Since it is common that genes participate in more than one pathway and drugs often affect to more than one gene, it is not infrequent that the predicted drug effects are accompanied of unexpected results. This fact reinforces the utility of comprehensive holistic modeling approaches like the one presented here. The gene intervention strategy implemented here is similar to the one used in the PathAct web tool [30] in the context of signaling pathway genes.

Obviously, off-target effects not described in DrugBank cannot be included in the predictions. However, Metabolizer would allow conjecturing new off-target effects by checking inconsistencies between the expected metabolic module activities from the prediction and the real ones observed upon the application of the drug.

### Automatic detection of optimal therapeutic targets

The *KnockOut* option of Metabolizer implements the *Auto Knockout* functionality to find the optimal KO to revert a condition. Within this modeling framework a gene KO is easy to simulate. Simply, the expression value is multiplied by 0.01. The model recalculates the module activity profiles. It is worth noting that a gene can participate in more than one module and that, depending on the location of the KO gene in the topology of the module, the KO can have a drastic or an irrelevant effect on the module activity.

Then, if two groups of samples are provided, metabolizer finds the KO intervention that makes samples of one of the classes resemble more to samples of the other class at the level of metabolic module activity profiles. This functionality has been designed to compare diseased to healthy conditions, or similar scenarios, and find the KO intervention that produces the maximum reversion from the disease to the healthy condition.

Firstly, a class predictor is built, using RF, that best discriminate among the two classes compared. Since only 553 genes participate in the modules, for each sample all the possible gene KOs can be carried out. For each simulated KO, the metabolic values are recalculated and the predictor estimates the possibility that the resulting metabolic profile belongs to the opposite class. All metabolic profiles resulting from the KO are ranked by this probability and the higher probabilities represent the most promising KO interventions. Combinations of KOs are not feasible in interactive mode but they can be experimented manually in the individual sample mode. Additional Figure 2 shows a schema of the procedure.

**Figure 2.**
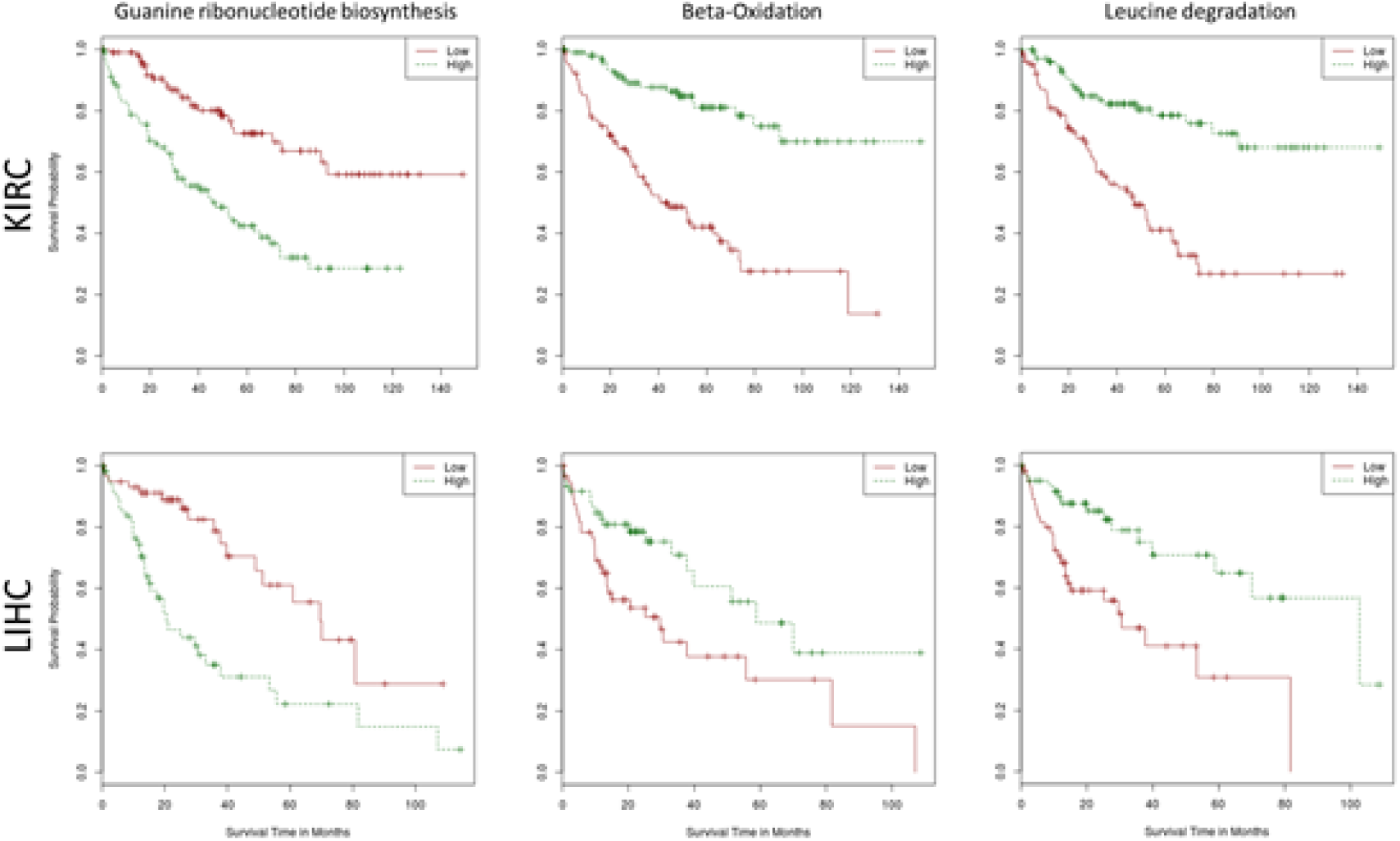
Kaplan-Meyer survival plots for the Guanine ribonucleotide biosynthesis module (left), the Beta-Oxidation module (center) and the Leucine degradation module (right) in KIRC (upper row) and LIHC (lower row) tumors.

### Implementation in a web server

Metabolizer is a web-based application that implements the above described functionalities. Metabolizer web client has been developed in HTML5 with web components while the server component is written in R programming language. The program recodes gene expression data (either from microarray or from RNA-seq) into estimates of enzymatic activities along the sequence of reactions that transform simple into complex metabolites or vice-versa. Metabolizer can be used for several purposes that include: 1) estimation of differential metabolic activity by comparing two conditions, 2) derivation of class predictors for further classification of new samples using metabolic activities as multigenic biomarkers; 3) search of therapeutic targets by predicting the ultimate impact of KOs on the final metabolite production activity of the modules, and 4) automatic detection of the optimal KO that makes the metabolic profile of an initial condition as close as possible to a final condition (e.g. the KO that reverts a disease to the normal status).

In addition to human metabolism, Metabolizer includes the metabolism of 5 more model species, namely Mouse, Rat, Zebrafish, Drosophila and Worm, taken from the KEGG repository too.

The input for Metabolizer consists of files of normalized gene expression values (in TSV format) along with an accompanying text file containing the experimental design. A tutorial explains in detail the required format.

The results produced include a graphical output that represents the metabolic modules analyzed in which the sequences of enzymatic reactions that transform simple into complex metabolites are highlighted. In this way, disruptions or activations in the metabolite transformation chain can be easily visualized providing a straightforward interpretation of its real impact on the ultimate metabolite production activity. A convenient graphic interface, based on the CellMaps [31] libraries, provides an interactive view of the metabolic modules with configurable color-coded representation of the metabolic modules and their components. In this interface, gene activity and module activities are simultaneously represented providing a visual, intuitive indication on relevant changes in the activity of the genes and their final impact in the activity of the modules (see Figure 1)

In addition, tables listing the modules showing a significant change in the activity are provided, along with the statistics and the corresponding p-values.

In addition to anonymous use for occasional users, Metabolizer allows user registration. In this case, all the comparisons and operations carried out are maintained in a user account.

Metabolizer can be found at: http://metabolizer.babelomics.org.

## Performance of the method

### Samples and data processing

We downloaded RNA-seq counts for a total of 2256 cancer samples and 285 healthy reference tissue samples, corresponding to Breast Invasive Carcinoma (BRCA), Liver Hepatocellular Carcinoma (LIHC), Kidney Renal Clear Cell Carcinoma (KIRC) and Prostate Adenocarcinoma (PRAD) cancer types (see Table 1), from The Cancer Genome Atlas (TCGA) repository (https://tcga-data.nci.nih.gov/tcga/). We used the COMBAT method [32] for batch effect correction and the trimmed mean of M-values normalization method (TMM) [33] for gene expression normalization. Normalized gene expression values were log-transformed and re-scaled between 0 and 1.

### Sensitivity and specificity of models of metabolic module activity

Using module activities estimated for all the samples a predictor able to differentiate cancer from healthy samples was built using RF [27] to estimate the sensitivity and specificity of modules to predict class membership. Since the number of features is not too high (there are only 95 modules), feature selection was not considered necessary here. Specifically, we repeated 50 times the 5-fold cross-validation on the dataset: two groups, one composed of normal samples and another, with the same size, composed of tumor samples randomly sampled were constructed. Four fifth parts were used to train a RF [27] predictor and the remainder fifth part was used to test the predictor with all the module activities. Since the real labels of the fifth part are known, the correct and wrong assignations per class were used to calculate the area under the ROC curve (AUC). Table 2 shows that the predictive power for the four cancer types in Table 1 using module activities as features is extremely high. The AUC in real class comparisons can be compared to the poor AUC values in artificial classes obtained by random permutation of cancer and control labels. This strongly suggests that module activities account for real biological features that change between cancers and normal tissues.

**Table 2.** AUC values obtained for tumor types in Table 1, with the corresponding AUC values obtained when artificial classes are obtained by randomizing sample labels.

|  | **BRCA** | **BRCA**<br>**random** | **LIHC** | **LIHC**<br>**random** | **KIRC** | **KIRC**<br>**random** | **PRAD** | **PRAD**<br>**random** |
| --- | --- | --- | --- | --- | --- | --- | --- | --- |
| **Mean** | 1.000 | 0.495 | 1.000 | 0.525 | 0.999 | 0.477 | 0.998 | 0.491 |
| **Standard Deviation** | 0.000 | 0.190 | 0.000 | 0.251 | 0.002 | 0.216 | 0.006 | 0.208 |
| **Median** | 1.000 | 0.5025 | 1.000 | 0.533 | 1.000 | 0.464 | 1.000 | 0.469 |
| **Median Absolute Deviation** | 0.000 | 0.205 | 0.000 | 0.312 | 0.000 | 0.243 | 0.000 | 0.228 |

### Comparison to other methods

Different approaches for the detection of different aspects of metabolic module activity have been proposed. In order to compare the accuracy of Metabolizer in detecting metabolic module activity, we have used a version of GSEA based on logistic regression [34] as implemented in the *mdgsa* Bioconductor package (http://bioconductor.org/packages/release/bioc/html/mdgsa.html) and a popular PT-based algorithm SPIA [35], as implemented in the *SPIA* Bioconductor package (http://bioconductor.org/packages/release/bioc/html/SPIA.html). For these methods, gene sets were defined using the genes within the metabolic modules. Additionally, the SPIA method requires also of the topology of the modules. In order to adapt the modules to the pathway format needed for the SPIA function the relations between metabolites on a module are considered as activations. GSEA detects only differential activity while SPIA and Metabolizer also detect whether this different activity implicates activation or deactivation. Four cancers (Table 1) were used for the comparison. The sensitivity of the method was measured as to the number of modules detected as differentially active by comparing the four cancers in Table 1 with respect to their corresponding healthy tissues. The specificity was measured as the number of differentially active modules (false positives) found by each method in a comparison involving individuals of the same class.

In addition, we utilized a well-known version of CBM method [36], as implemented in the IMAT tool [37] using the human metabolic network Recon 2 V2.02 [38], for comparing its performance to Metabolizer as well. The IMAT tool maximizes the number of highly expressed reactions that are active and the number of lowly expressed reactions that are inactive. The reaction activity is inferred from the binarization of the corresponding gene expression values following a Boolean logic from Gene-Protein-Reaction (GPR) rules within the context of metabolic networks [39]. *CPLEX* (V12.6.2) solver was used for solving *Mixed Integer Linear Programming* problems. Optimum solutions provide flux values of reactions and these flux values were used to classify reactions as active and inactive. All parameters were set as in the original article [36]. The binary results of reactions (active/inactive) were used to train a classifier. Since this CBM method is based on a pathway definition (Recon 2) [38] which is different from the KEGG metabolic modules used here [18], we use a different benchmarking framework in which reaction values are used as predictor features [40].

Given that classifiers based either on Module activities or on CBM reaction activities were able of distinguishing between cancer and normal tissues with almost 100% accuracy we challenged them with a more complex classification problem: distinguishing between cancer subtypes in the case of breast cancer. The BRCA dataset (Table 1) contains PAM50-defined [41] subtypes Basal-like, HER2-enriched, Luminal A and Luminal B of Breast Invasive Carcinoma [42]. The performance of a RF [27] classifier trained using Metabolizer module activities and reaction activities obtained by CBM were compared by 5-fold cross validation, using gene expression based classification as a gold standard. It is worth noticing that only one gene belonging to the metabolic modules, *PHGDH*, was in the list of PAM50 genes used to define the BRCA subtypes.

We first compared the capability for detecting differentially activated modules when cancer is compared to the corresponding unaffected tissue in four distinct cancer types: BRCA, LIHC, KIRC and PRAD (Table 1). For this contrast, we used the conventional approach based on unstructured gene sets, the GSEA [34], and an approach that takes into account the relationships among genes within gene sets, the SPIA [35]. Table 3 shows the number of modules found as differentially activated in the different cancers by the different methods. Metabolizer outperforms both the sensitivity and specificity of GSEA and SPIA. GSEA founds between 5 and 14 modules, depending on the cancer, with averages ranging from 2 to 7 false positive (FP) modules (about 50% of false discovery). SPIA increases the specificity at the exchange of reducing the sensitivity, with a very low detection rate. Metabolizer increases by almost one order of magnitude both sensitivity and specificity (Table 3).

**Table 3.**
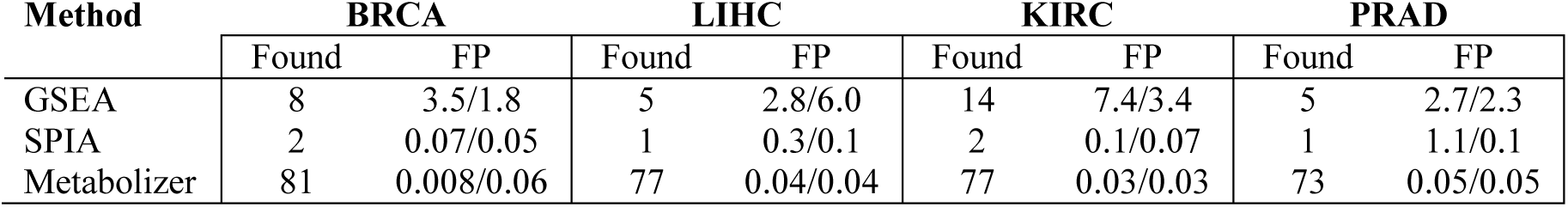
Number of modules found as differentially activated in the cancers listed in Table 1 by the different methods GSEA, SPIA and Metabolizer. The number of false positives (FP) was calculated by comparing 1000 times two artificial sample sets by random sampling of normal tissues maintaining the proportions of the real comparison. That is 102 vs 11 for BRCA, 41 vs 7 in LIHC, 63 vs 9 in KIRC and 46 vs 6 in PRAD. The same procedure was repeated using cancer samples. In this case the proportions were 995 vs 102 in BRCA, 253 vs 41 in LIHC, 463 vs 63 in KIRC and 333 vs 46 in PRAD. The second column for each cancer type shows the average number of FPs obtained with normal samples / the same figure obtained from cancer samples.

| Method | BRCA |  | LIHC |  | KIRC |  | PRAD |  |
| --- | --- | --- | --- | --- | --- | --- | --- | --- |
|  | Found | FP | Found | FP | Found | FP | Found | FP |
| GSEA | 8 | 3.5/1.8 | 5 | 2.8/6.0 | 14 | 7.4/3.4 | 5 | 2.7/2.3 |
| SPIA | 2 | 0.07/0.05 | 1 | 0.3/0.1 | 2 | 0.1/0.07 | 1 | 1.1/0.1 |
| Metabolizer | 81 | 0.008/0.06 | 77 | 0.04/0.04 | 77 | 0.03/0.03 | 73 | 0.05/0.05 |

Additional Table 2 contains the modules detected as differentially active by these three methods. In general, the results found by the methods were consistent across them, taking into account their different sensitivities. As expected, modules controlling the biosynthesis of nucleotide precursors [43] and Acetyl-CoA [44,45] were found across cancers by GSEA and Metabolizer. However, several well-known metabolic activities associated to cancer development and progression, such as increased production of L-Proline [46] and succinate [47], or related to metastasis, such as Fumarate [48], 4-Aminobutanoate (GABA biosynthesis) [49] or N-Acylsphingosine (Ceramide biosynthesis) [50], were found only by the more sensitive Metabolizer method.

Since CBM analysis is based on a different type of pathway (Recon 2), the comparison cannot be carried out in the previous benchmarking framework that uses metabolic modules defined within KEGG pathways. Instead, we carried out a comparison of classification performances using a previously proposed benchmarking framework based on the use of reaction activities estimated by CBM as features for classification [40]. Given that cancer vs normal tissue was a quite naive classificatory problem for which both CBM and Metabolizer resulted in almost 100% classification accuracy, we used a more challenging classification problem: BRCA subtype prediction. Classification performances were carried out using a RF predictor with 5-fold cross validation. Since BRCA subtypes have been defined using the expression of 50 genes with the PAM50 classifier [41], the classification obtained using the expression of all genes is expected to provide an upper limit of classification performance. Figure 3 shows how module activities obtained with Metabolizer outperform CBM-based reaction activities in classifying all the BRCA subtypes.

**Figure 3.**
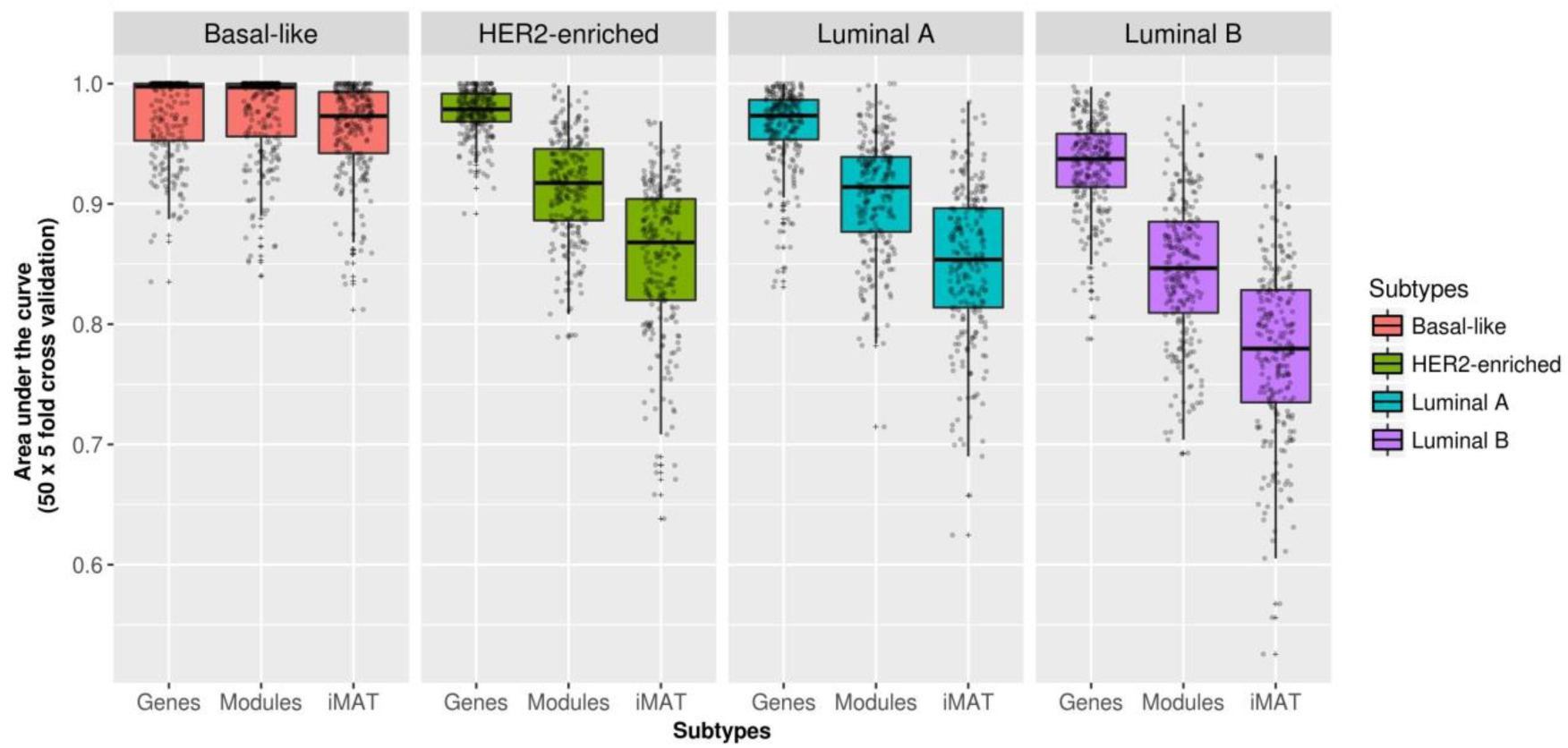
Classification performance obtained using module activities inferred with Metabolizer and CBM-based reaction activities for the prediction of BRCA subtypes. BRCA subtypes are defined on the bases of PAM50 gene activities and therefore, gene expression is taken as the gold standard classification performance.

## Validation and case uses

### An example of automatic optimal KO

To illustrate the potential of the auto-KO option we have used this tool to find KOs that would make a KIRC sample as similar as possible to a normal kidney sample in terms of metabolism. We used a balanced dataset composed of the 72 normal kidney samples available and 72 KIRC samples randomly sampled among all the available tumor patients and used the Auto-KO option. Then, a class predictor is built that will be used to decide to what extent the tumor sample, after the KO, could be identified as a normal sample. Figure 4 shows how most of the KOs does not have an effect that revert the metabolic tumor status towards a normal kidney that increases the probability of being recognized as normal by the predictor. However, in a few cases the result of the KO changes the metabolic status of the tumor in a way that is identified as normal in approximately a 25% (those in the small red peak approximately in 0.25). Table 4 lists the genes whose KO changes the metabolism of tumor cell and makes it more “normal”.

**Figure 4.**
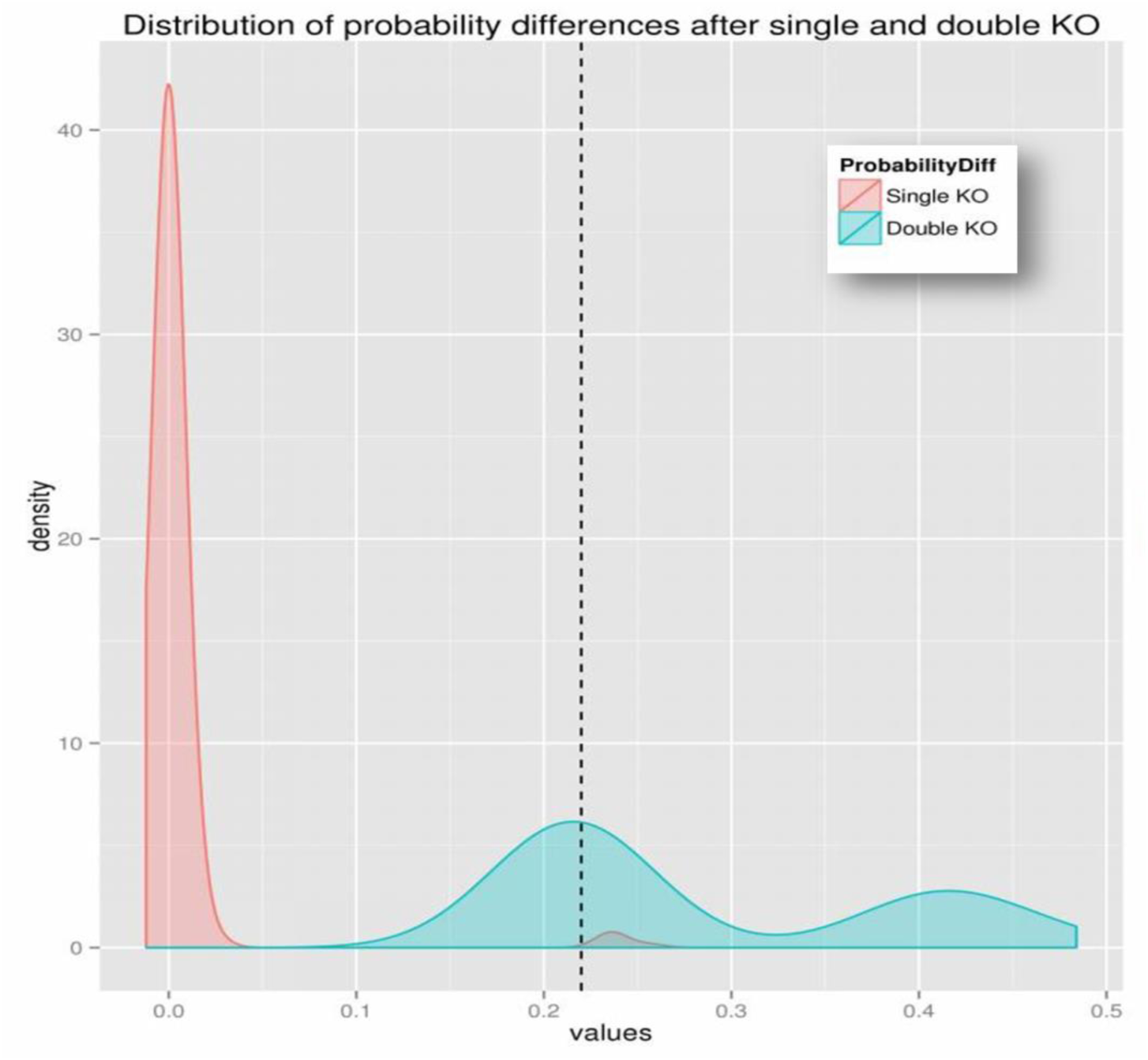
Distribution of the difference of probabilities that the predictor identifies a sample as a normal cell after and before the KOs of the corresponding genes (red distribution) or pair of genes (blue distribution).

**Table 4.** Probabilities of being identified as normal cells after the KO of the gene.

| Gene Symbol | Entrez ID | P(normal)<br>after KO | P(normal)<br>before KO | Change in<br>probability |
| --- | --- | --- | --- | --- |
| HSD17B12 | 51144 | 0.348 | 0.092 | 0.256 |
| TECR | 9524 | 0.348 | 0.092 | 0.256 |
| SC5D | 6309 | 0.328 | 0.092 | 0.236 |
| EBP | 10682 | 0.328 | 0.092 | 0.236 |
| DHCR24 | 1718 | 0.328 | 0.092 | 0.236 |
| LSS | 4047 | 0.328 | 0.092 | 0.236 |
| TM7SF2 | 7108 | 0.328 | 0.092 | 0.236 |
| NSDHL | 50814 | 0.328 | 0.092 | 0.236 |
| CYP51A1 | 1595 | 0.328 | 0.092 | 0.236 |
| HSD17B7 | 51478 | 0.328 | 0.092 | 0.236 |
| DHCR7 | 1717 | 0.328 | 0.092 | 0.236 |

### Validation of optimal KO predictions

Some of the optimal KO predictions were known cancer-related genes. For example, *HSD17B12*, is a known cancer antigen [51], *EBP* is a long known cancer estrogen receptor [52] or *DHCR24* is a gene whose over-expression is related to bad prognostic in several cancers [53], which explain the potential predicted impact that their KOs have in the cancer metabolic profile.

However, beyond the knowledge derived from the literature, other experimental evidences, such as the recent release of a large-scale map of cancer dependency [54], can be used to validate predictions made on the simulated KOs that would potentially reduce the cancer phenotype of cells and make them resemble normal cells. The expectation is that inhibitions of optimal KO genes should result in the reduction of the proliferative capability of the corresponding cell lines that could be interpreted as a reversion of cancer phenotype towards a normal cell (or at less, a non-proliferative cell). In spite of the fact that cancer outcome is a much more complex phenotype than the proliferation of a cell line, when genes in Table 4 are inhibited in the cancer dependency experiment [54] a reduction in the proliferation was observed for ten out of the eleven predicted optimal KOs (*HSD17B12, TECR, SC5D, EBP, DHCR24, LSS, NSDHL, CYP51A1, HSD17B7, DHCR7*) (see Figure 5). Moreover, in some cases we were able to detect an increase of patient survival in patients with low expression of some of the optimal KO proteins in Table 4 (assuming that low cancer cell survival represents the same effect that patient survival). Thus, according to Protein Atlas, low expression of *TECR* protein is significantly associated to better patient survival in urothelial cancer (see https://www.proteinatlas.org/ENSG00000099797-TECR/pathology/tissue/urothelial+cancer), and the same is observed in *DHCR24* in endometrial cancer (https://www.proteinatlas.org/ENSG00000116133-DHCR24/pathology/tissue/endometrial+cancer), *LSS* in urothelial cancer (https://www.proteinatlas.org/ENSG00000160285-LSS/pathology/tissue/urothelial+cancer), *CYP51A1* in cervical cancer (https://www.proteinatlas.org/ENSG00000001630-CYP51A1/pathology/tissue/cervical+cancer), and *HSD17B7* in renal cancer (https://www.proteinatlas.org/ENSG00000132196-HSD17B7/pathology/tissue/renal+cancer).

**Figure 5.**
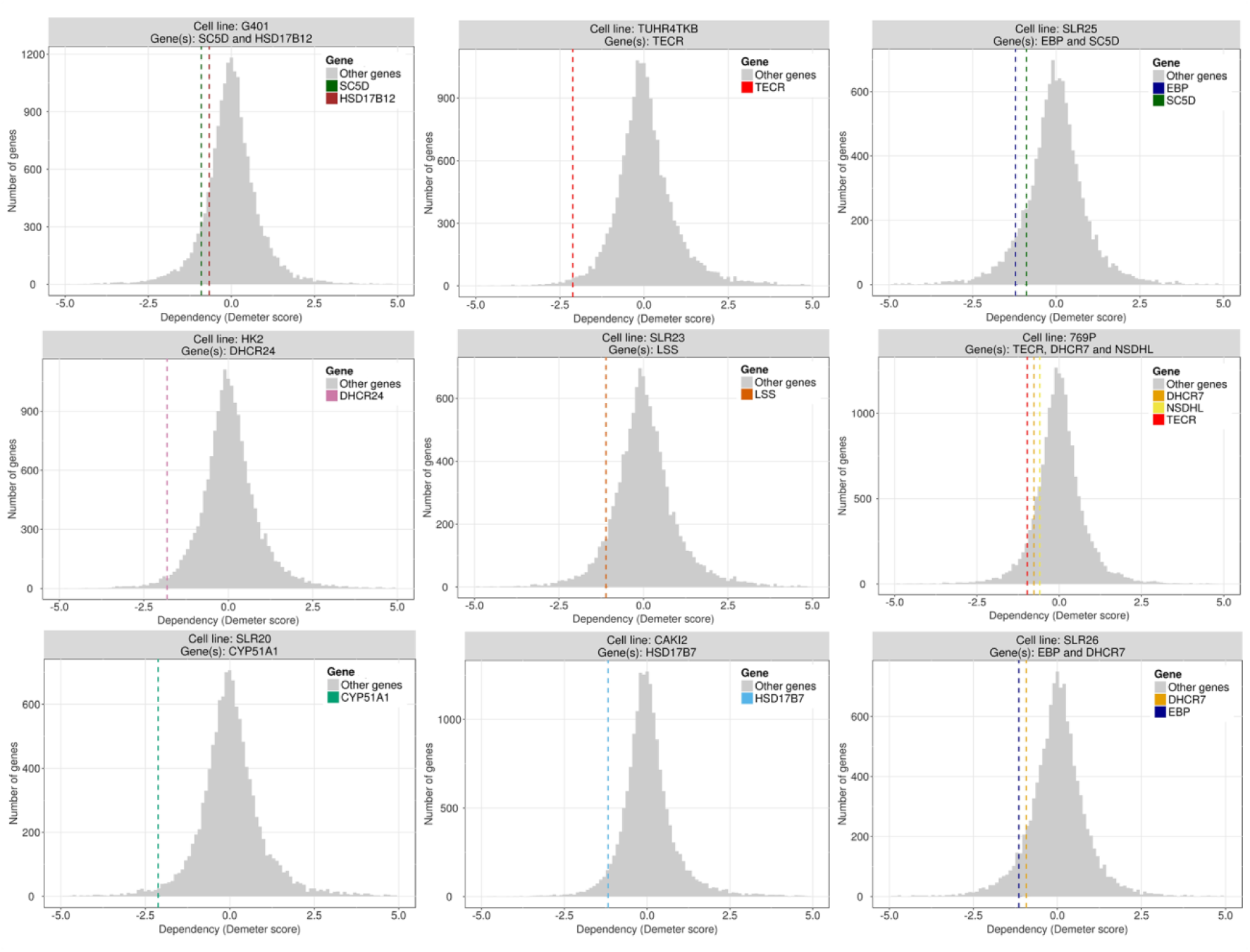
Essentiality (Demeter score) of genes predicted as optimal KOs with respect to the background distribution of essentiality values. Values below 0 indicate lower proliferation. From left to right and top to bottom: *HSD17B12* and *SC5D* in cell line G401 (KIDNEY); *TECR* in cell line TUHR4TKB (KIDNEY) (this gene shows the same results in KMRC1 cell line of KIDNEY, data not shown); *SC5D* and *EBP* in SLR25 cell line (KIDNEY) (*SC5D* shows the same result in G401 cell line of SOFT_TISSUE, data not shown); *DHCR24* in cell line HK2 (KIDNEY); *LSS* in cell line SLR23 (KIDNEY); *NSDHL, DHCR7* and TECR in 769P cell line (KIDNEY); *CYP51A1* in cell line SKRC20 (KIDNEY) (also less proliferative in SLR20 KIDNEY cell line, data not shown); *HSD17B7* in cell line CAKI2 (KIDNEY); *DHCR7* and *EBP* in cell line SLR26 (KIDNEY).

### Experimental validation of an optimal KO prediction

Finally, as an additional validation, we used the optimal KO option of Metabolizer in a different cancer type, gastric cancer patients (STAD). Table 5 shows the predictions. The gene causing the strongest effect, *DPYS*, was found as essential in the catalog of cancer dependencies [54]. The second predicted gene, *UPB1*, encodes an enzyme (β-ureidopropionase) that catalyzes the last step in the pyrimidine degradation pathway, required for epithelial-mesenchymal transition [55]. Using a cancer model of gastric adenocarcinoma (AGS cell line) we carried out a cell proliferation experiment upon depletion of *UPB1* gene expression. The shRNAs targeting *UPB1* were purchased from the MISSION (Sigma Aldrich) library, catalog SHCLNG-NM_016327. Lentivirus production and transduction was performed following standard protocols and cell cultures were selected with puromycin for 72 hours prior cell seeding for evaluation of proliferation/viability by methylthiazol tetrazolium (MTT)-based assays (Sigma-Aldrich). The data corresponds to sextuplicates and was replicated in different assays. *UPB1* expression was detected with the Human Protein Atlas HPA000728 antibody (Sigma-Aldrich) and gene expression measured with primers 5’-TCGACCTAAACCTCTGCCAG-3’ and 5’-TAAGCCTGCCACACTTGCTA-3’, using PPP1CA as control. As anticipated by our prediction, three different short hairpin shRNA sequences directed to *UPB1* caused a significant decrease in cell proliferation (see Figure 6). This result constitutes an independent validation that reinforces the prediction made by the model proposed.

**Figure 6.**
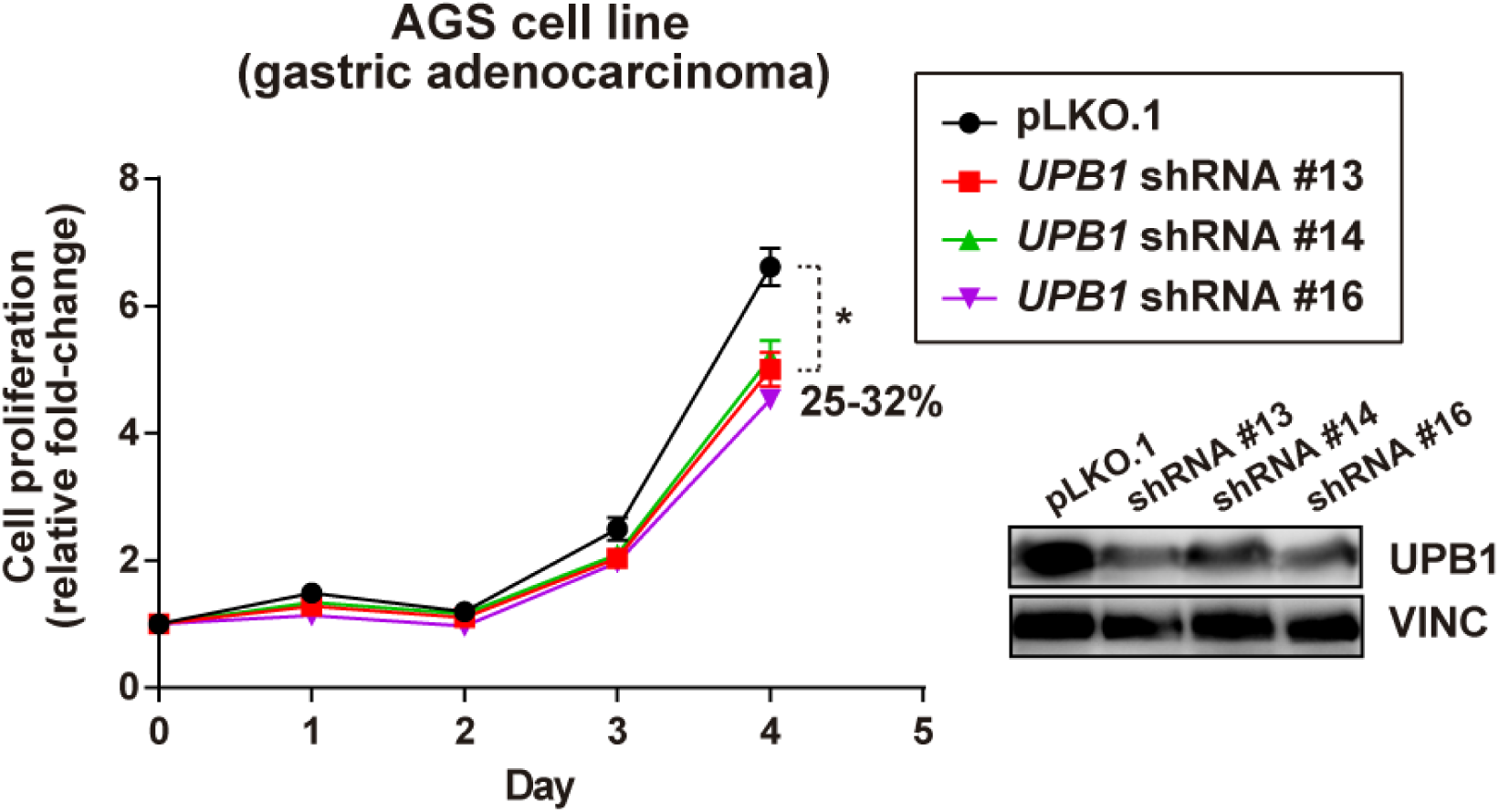
Relative cell proliferation of line AGS (stomach gastric adenocarcinoma) upon *UPB1* expression depletion by three different MISSION shRNAs or transduced with control vector pLKO.1. The asterisk indicates significant differences (Mann-Whitney test p-values < 0.01). The percentage of reduction of cell proliferation is also shown. The prediction of *UPB1* essentiality made by Metabolizer was confirmed by a relatively more sensitive behavior.

**Table 5.**
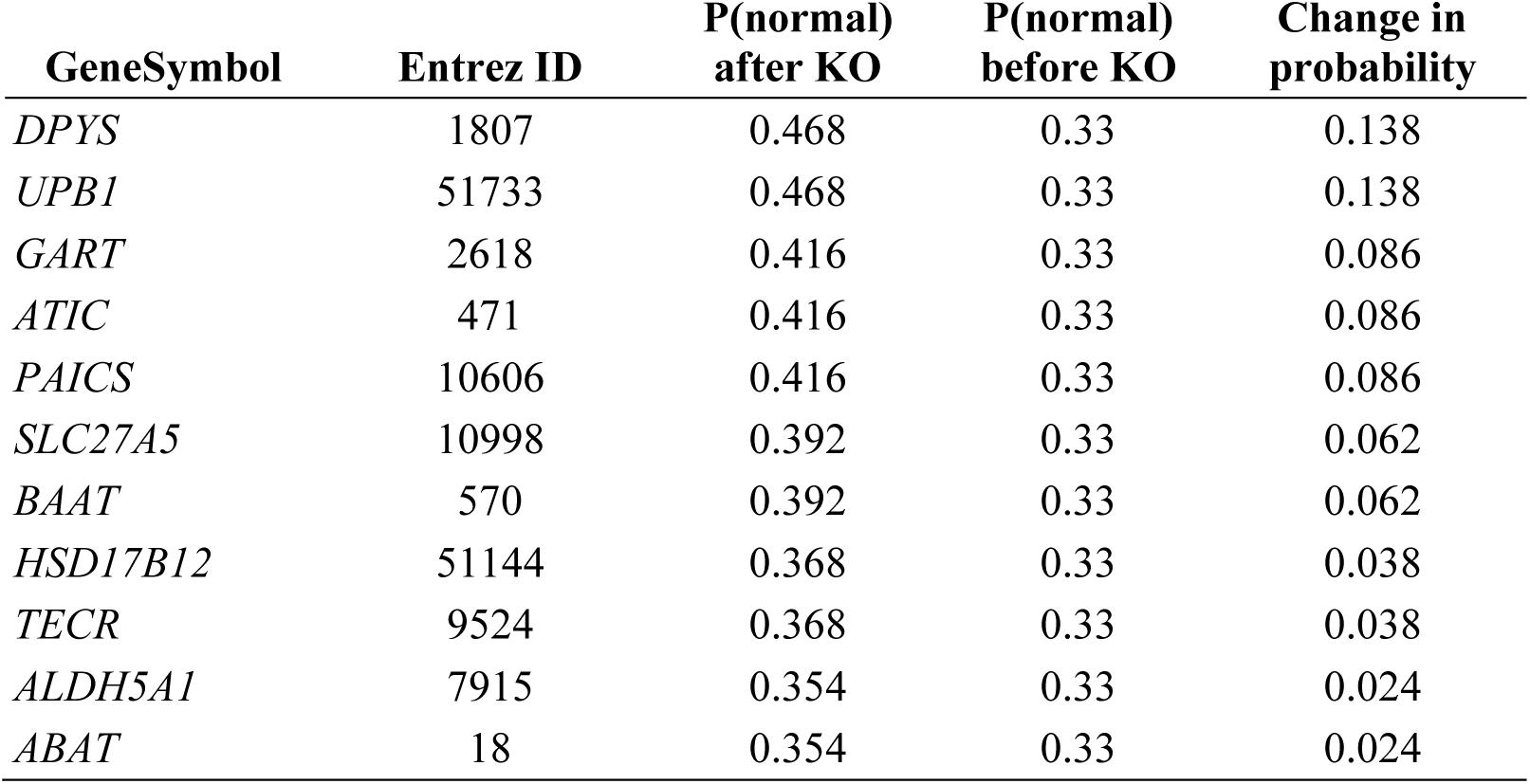
Probabilities of being identified as normal cells after the KO of the gene.

All these different kind of observations strongly support the validity of the predictions made.

### Module activities associated to patient survival

In order to know whether the modeled activity of metabolic modules account for a phenotype as complex as cancer prognostic, we have used gene expression data along with survival data corresponding to two different cancers, Kidney renal clear cell carcinoma (KIRC) and Liver hepatocellular carcinoma (LIHC). These cancers were selected because they have a balanced number of patients alive and deceased, which allows estimations of Kaplan-Meier (K-M) survival curves [56]. Survival data were obtained from the cBIOportal [57].

The value of the activity estimated for each module in each individual was used to assess its relationship with individual patient survival using K-M curves [56]. Calculations were carried out using the function *survdiff* from the *survival* R package [58], which tests for significant differences when two groups of patients are compared. Then, for each module, survival analysis comparing patients with high module activity to patients with low module activity was carried out. The high module activity and low module activity groups were defined by patients within the 20% upper and lower percentiles of module activity, respectively. The results were adjusted for multiple testing effects using the FDR method [26]. Table 6 shows the modules whose activity was found to be significantly associated to patient survival in some of the cancers. Despite a comprehensive description of the results is beyond the scope of this manuscript, it is worth mentioning how the production of nucleotides and precursors (GTP, UMP and CDP) shows a recurrent significant activation in both cancers. Genes in the corresponding modules are targeted by well-known anticancer clinical drugs, such as Gemcitabine, which is approved for the treatment of at least four advanced cancer types, and Mercaptopurine (DB00441 and DB01033 entries in DrugBank), respectively. The mechanism of action of these drugs is based on the inhibition of DNA synthesis that leads to cell death by specifically inhibiting the production process of GTP, CDP and their precursor metabolites. Also, *Fatty acid biosynthesis, Inositol phosphate metabolism* and *Beta-Oxidation* modules are highly correlated with patient survival. Actually, activation of de novo fatty acids synthesis, is exclusive of cancer cells and has an essential role in supporting conversion of nutrients into metabolic components for membrane biosynthesis, energy storage and generation of signaling molecules [59]. In fact, several preclinical and clinical studies have been addressed to test the effect of inhibiting fatty acid synthase (*FASN*) in different cancer types. It is also known the use of serine by many cancers [60] which explains the significant association to survival found for the *Serine biosynthesis* module. Figure 2 shows the Kaplan-Meyer survival plots for the Guanine ribonucleotide biosynthesis, the Beta-Oxidation and the Leucine degradation modules, corresponding to the main metabolic processes of nucleotides, amino acids and lipids, in KIRC and LIHC tumors.

**Table 6.**
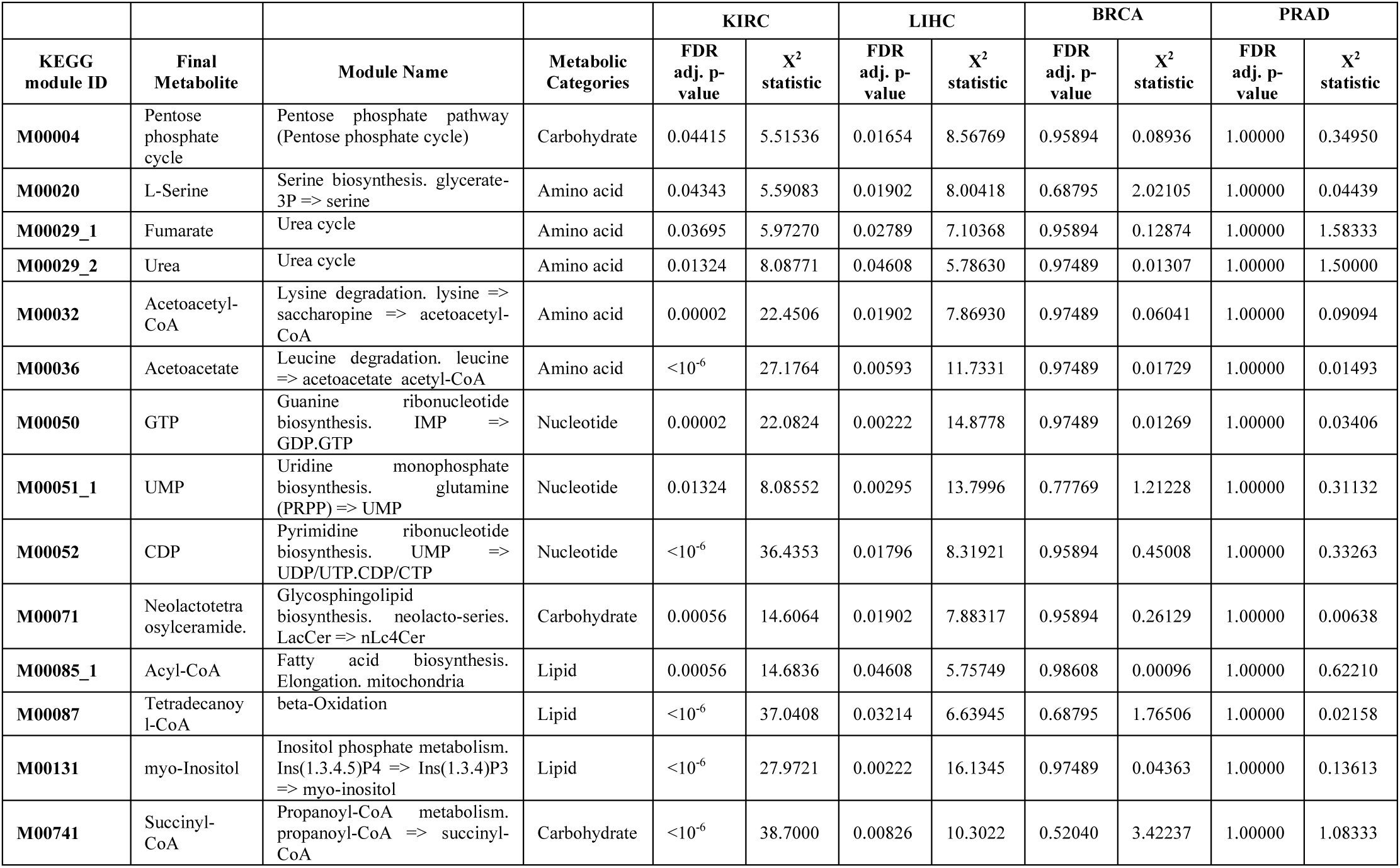
KEGG modules with activity significantly associated to patient survival in both KIRC and LIHC tumors. BRCA and PRAD did not show any significant result.

The rest of modules are related to the production of metabolites involved in cell proliferation or other processes connected to cancer origin or progression.

### Synergistic effects

Although a 25% increase in the probability of resembling a normal cell might look a small value, it is not expectable that a unique intervention restores a cancer cell to a normal cell status. However, if we assay combinations of double KOs between genes in Table 4, we observe a dramatic increase in the “normal” character of the cancer cell, as graphically depicted in Figure 4. Most of the combinations have similar effect that the single KOs alone (peak around 0.25). However, some combinations produce synergistic changes with a dramatic effect in the metabolic profile of KIRC cells that make them more similar to normal cells than to cancer cells (peak around a 0.5 of change in the probability). Table 7 shows how the probability of being identified as normal cells for 17 combinations is higher than 0.5.

**Table 7.** Probabilities of being identified as normal cells after double KOs of the genes

| Gene1 | Gene2 | P(normal)<br>after KO | P(normal)<br>before KO | Change in<br>probability |
| --- | --- | --- | --- | --- |
| TECR | LSS | 0.576 | 0.092 | 0.484 |
| TECR | DHCR24 | 0.560 | 0.092 | 0.468 |
| TECR | EBP | 0.538 | 0.092 | 0.446 |
| TECR | CYP51A1 | 0.542 | 0.092 | 0.450 |
| HSD17B12 | EBP | 0.530 | 0.092 | 0.438 |
| HSD17B12 | DHCR7 | 0.524 | 0.092 | 0.432 |
| TECR | SC5D | 0.554 | 0.092 | 0.462 |
| TECR | HSD17B7 | 0.526 | 0.092 | 0.434 |
| HSD17B12 | DHCR24 | 0.508 | 0.092 | 0.416 |
| HSD17B12 | CYP51A1 | 0.522 | 0.092 | 0.430 |
| HSD17B12 | HSD17B7 | 0.514 | 0.092 | 0.422 |
| TECR | NSDHL | 0.536 | 0.092 | 0.444 |
| TECR | DHCR7 | 0.526 | 0.092 | 0.434 |
| HSD17B12 | TM7SF2 | 0.508 | 0.092 | 0.416 |
| HSD17B12 | NSDHL | 0.502 | 0.092 | 0.410 |
| HSD17B12 | SC5D | 0.508 | 0.092 | 0.416 |
| TECR | TM7SF2 | 0.510 | 0.092 | 0.418 |
| HSD17B12 | LSS | 0.486 | 0.092 | 0.394 |
| SC5D | LSS | 0.346 | 0.092 | 0.254 |
| TM7SF2 | HSD17B7 | 0.356 | 0.092 | 0.264 |
| DHCR24 | LSS | 0.340 | 0.092 | 0.248 |
| DHCR24 | TM7SF2 | 0.326 | 0.092 | 0.234 |
| EBP | HSD17B7 | 0.346 | 0.092 | 0.254 |
| HSD17B7 | DHCR7 | 0.350 | 0.092 | 0.258 |
| SC5D | DHCR24 | 0.324 | 0.092 | 0.232 |
| TM7SF2 | DHCR7 | 0.316 | 0.092 | 0.224 |
| DHCR24 | CYP51A1 | 0.330 | 0.092 | 0.238 |
| SC5D | EBP | 0.326 | 0.092 | 0.234 |
| SC5D | DHCR7 | 0.332 | 0.092 | 0.240 |
| CYP51A1 | DHCR7 | 0.336 | 0.092 | 0.244 |
| CYP51A1 | HSD17B7 | 0.306 | 0.092 | 0.214 |
| SC5D | CYP51A1 | 0.312 | 0.092 | 0.220 |
| DHCR24 | DHCR7 | 0.336 | 0.092 | 0.244 |
| NSDHL | CYP51A1 | 0.330 | 0.092 | 0.238 |
| LSS | CYP51A1 | 0.334 | 0.092 | 0.242 |
| EBP | NSDHL | 0.312 | 0.092 | 0.220 |
| EBP | CYP51A1 | 0.318 | 0.092 | 0.226 |
| SC5D | NSDHL | 0.310 | 0.092 | 0.218 |
| EBP | LSS | 0.328 | 0.092 | 0.236 |
| DHCR24 | HSD17B7 | 0.292 | 0.092 | 0.200 |
| LSS | TM7SF2 | 0.340 | 0.092 | 0.248 |
| LSS | NSDHL | 0.324 | 0.092 | 0.232 |
| NSDHL | DHCR7 | 0.306 | 0.092 | 0.214 |
| TM7SF2 | NSDHL | 0.328 | 0.092 | 0.236 |
| TM7SF2 | CYP51A1 | 0.328 | 0.092 | 0.236 |
| SC5D | TM7SF2 | 0.314 | 0.092 | 0.222 |
| HSD17B12 | TECR | 0.312 | 0.092 | 0.220 |
| EBP | DHCR24 | 0.304 | 0.092 | 0.212 |
| EBP | TM7SF2 | 0.306 | 0.092 | 0.214 |
| LSS | DHCR7 | 0.324 | 0.092 | 0.232 |
| SC5D | HSD17B7 | 0.304 | 0.092 | 0.212 |
| LSS | HSD17B7 | 0.320 | 0.092 | 0.228 |
| NSDHL | HSD17B7 | 0.268 | 0.092 | 0.176 |
| DHCR24 | NSDHL | 0.310 | 0.092 | 0.218 |
| EBP | DHCR7 | 0.304 | 0.092 | 0.212 |

As an example, Figure 7 shows the double KO in genes *SC5D* and *TERC*, which affects to the *Fatty acid elongation* module and the *Steroid biosynthesis* module, the last one involved in the production of cholesterol. Actually, it is long known that tumor membranes are rich in cholesterol [61], suggesting that cholesterol utilization by cancer cells is an important feature of carcinogenesis and, probably, metastasis [62].

**Figure 7.**
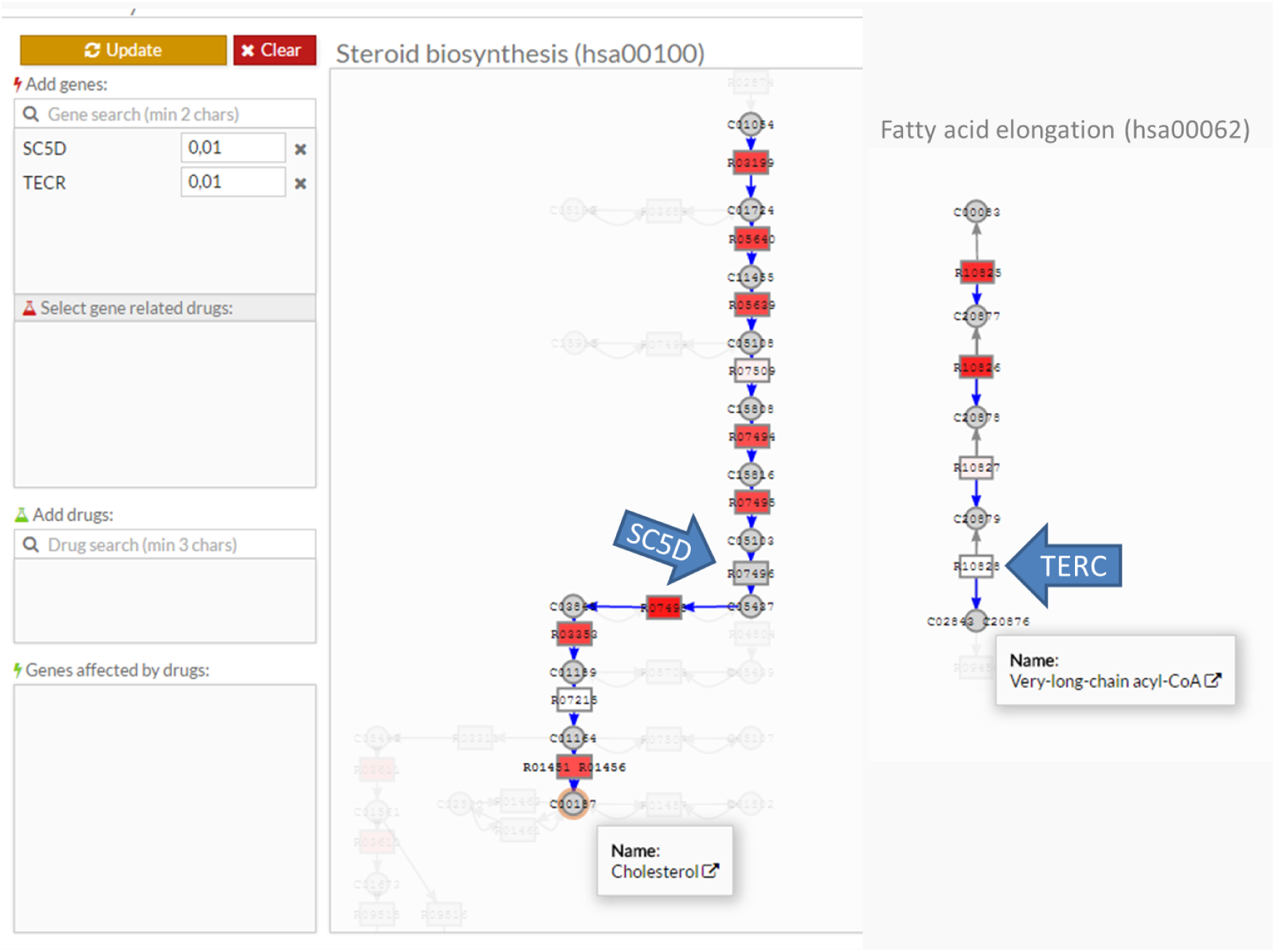
Representation of the modules corresponding to the Steroid biosynthesis (hsa00100) and Fatty acid elongation (hsa00062). Arrows point to the KO genes (SC5D and TERC). KOs were made by substituting the actual gene expression values by a low non-zero value of 0.01 (as can be seen in the Add genes box of the Metabolizer program.) Genes in red indicate that they were active. At the end of both modules the resulting metabolite can be seen.

## Conclusions

The associations found between metabolic module activities and patient survival confirms that metabolic modules can be realistically modeled within the proposed framework. Moreover, metabolic module activities obtained under the proposed modeling method outperform other methods used to infer metabolic activity, such as GSEA [8], SPIA [35] or CBM [36] (as implemented in IMAT tool [37]). And, furthermore, we have validated most of the predictions made by the method in an independent dataset. These results show that metabolic modules can be considered a relevant type of functional module in cancer and probably also in other diseases related to metabolism. The program Metabolizer allows researchers to easily estimate module metabolic activities from gene expression measurements and use them for different purposes. Thus, the comparison between two conditions can throw light on the subjacent molecular mechanisms that make them different. In this way, disease mechanisms or drug mechanisms of action can easily be interpreted within the context of metabolism. Such comparisons can also be used to derive multigenic predictors with a mechanistic meaning, that have demonstrated to be useful to predict complex traits [4].

Diagnostic strategies are rapidly changing in cancer and other diseases because of the availability of increasingly affordable genomic analysis [63]. Therapies that specifically target genetic alterations are probing to be safer and more effective than traditional chemotherapies when used in the adequate patient population [64]. Perhaps, one of the most relevant aspects of modeling is that models allow predicting the effect of simulated gene expression profiles over the activity of metabolic modules, opening the door to anticipate the effect of intervention on genes. In this respect, Metabolizer constitutes an extremely useful tool for finding putative actionable targets for a specific condition [65]. This is very relevant in the context of personalized medicine and can help in finding individualized therapeutic interventions for patients [66]. In fact, recent reports indicate that genes involved in metabolic pathways show a remarkable heterogeneity across different cancer patients [67]. This suggests that personalized therapies might likely be successful providing the context of the interventions can be properly explored and understood with a tool such as Metabolizer. For example, synthetic lethality, defined as genetic mutations or gene expression alterations with little or null individual effect on cell viability but that results in cell death when combined, offers a promising range of potential therapeutic interventions [68] that can only be properly exploited in a framework such as the one provided by Metabolizer.

Therefore, Metabolizer can be considered an innovative tool that enables the use of standard measurements of gene expression in the context of the complexity of the metabolic network, with a direct application in clinic as well as in research in animal models.

## List of abbreviations

AUC: :Area Under the Curve
BRCA: :Breast Invasive Carcinoma
CBM: :Constraint Based Models
TSV: :Tab-Separated Values
FDR: :False Discovery Rate
GPR: :Gene-Protein-Reaction
GSEA: :Gene Set Enrichment Analysis
KEGG: :Kyoto Encyclopedia of Genes and Genomes
KGML: :KEGG Markup Language
KIRC: :Kidney Renal Clear Cell Carcinoma
K-M: :Kaplan-Meier
KO: :Knock-Out
LIHC: :Liver Hepatocellular Carcinoma
MoA: :Mode of Action
PRAD: :Prostate Adenocarcinoma
RF: :Random Forests
ROC: :Receiver Operating Characteristic
SVM: :Support Vector Machines
TCGA: :The Cancer Genome Atlas
TMM: :Trimmed Mean of M-values

## Declarations

Availability of data and materials

All the gene expression and clinical data used in this paper are available at the TCGA repository (https://tcga-data.nci.nih.gov/tcga/).

Metabolizer is available at: http://metabolizer.babelomics.org.

The source code is available at: https://github.com/babelomics/metabolizer

### Authors’ contributions

CC developed the propagation algorithm and the software to run it as well as analyzed the data, KR: helped with the development of the web interface, MRH: helped in the mathematical aspects of the propagation algorithm, FS: developed the web application tool, AA collaborated in the analysis of the data, MAP. FM and CH carried out the experimental validation, JCC helped in the development of the code used and JD conceived the work and wrote the paper.

## Additional Files

### Additional Table 1

Format: Excel file (.xlsx)

List of the modules used in this study. First column: Module name, second column: Main metabolic category, third column: Metabolic Sub-category of main category (if exist), fourth column: Description/Name of the module, fifth column: KGML identifier, sixth column: KEGG metabolic pathway in which the module is included, seventh column: URL to KEGG for the module, eight column: KEGG ID and name of the first metabolite in the module, ninth column: KEGG ID and name of the last metabolite in the module.

### Additional Table 2

Format: Excel file (.xlsx)

Modules with a significantly different activity between cancer and the corresponding healthy tissue in four different cancer types (BRCA, KIRC, LIHC and PRAD) detected by GSEA, SPIA and Metabolizer methods. Up/Down makes reference to activations/deactivations. Unknown appears in methods that only report a change in the activity but do not report the direction of this change. Value in parentheses correspond to the adjusted p-value.

### Additional Figure 1

Format: TIFF file (.tif)

Procedure used to estimate reaction node activity from the constituent gene expression activities. 1) If an enzyme is composed for more than one gene, the enzyme activity value is obtained as the 90^th^ percentile of normalized gene expression (to avoid possible outliers or artifacts). 2) When an isozyme is a complex, composed by more than one enzyme, the complex isozyme activity is obtained as the minimum activity value of all the involved enzymes (the limiting reaction capability). When a reaction node is composed by more than one isozyme, then, the maximum isoenzyme activity value is taken as the activity of the reaction node. 3) Finally, the module activity is inferred from the corresponding node activities.

### Additional Figure 2

Format: TIFF file (.tif)

*Auto Knockout* functionality to find the optimal KO to revert a condition. Initially, a predictor is trained to distinguish between normal and tumor samples. Then, for the tumor sample problem, all the possible KOs are simulated by sequentially multiplying by 0.01 the expression values of any of the genes involved in all the modules. Then, the theoretical KO profiles are calculated for each simulated KO sample and the predictor is used to assign a probability to belong to a normal or a tumor class. The rank of more likely belonging to the normal class is a rank of potential of transformation of a tumor sample into a normal sample.

